# Stable unmethylated DNA demarcates expressed genes and their cis-regulatory space in plant genomes

**DOI:** 10.1101/2020.05.21.109744

**Authors:** Peter A Crisp, Alexandre P Marand, Jaclyn M Noshay, Peng Zhou, Zefu Lu, Robert J Schmitz, Nathan M Springer

## Abstract

The genomic sequences of crops continue to be produced at a frenetic pace. However, it remains challenging to develop complete annotations of functional genes and regulatory elements in these genomes. Here, we explore the potential to use DNA methylation profiles to develop more complete annotations. Using leaf tissue in maize, we define ∼100,000 unmethylated regions (UMRs) that account for 5.8% of the genome; 33,375 UMRs are found greater than 2 kilobase pairs from genes. UMRs are highly stable in multiple vegetative tissues and they capture the vast majority of accessible chromatin regions from leaf tissue. However, many UMRs are not accessible in leaf (leaf-iUMRs) and these represent a set of genomic regions with potential to become accessible in specific cell types or developmental stages. Leaf-iUMRs often occur near genes that are expressed in other tissues and are enriched for transcription factor (TF) binding sites of TFs that are also not expressed in leaf tissue. The leaf-iUMRs exhibit unique chromatin modification patterns and are enriched for chromatin interactions with nearby genes. The total UMRs space in four additional monocots ranges from 80-120 megabases, which is remarkably similar considering the range in genome size of 271 megabases to 4.8 gigabases. In summary, based on the profile from a single tissue, DNA methylation signatures pinpoint both accessible regions and regions poised to become accessible or expressed in other tissues. UMRs provide powerful filters to distill large genomes down to the small fraction of putative functional genes and regulatory elements.

**Significance Statement:** Crop genomes can be very large with many repetitive elements and pseudogenes. Distilling a genome down to the relatively small fraction of regions that are functionally valuable for trait variation can be like looking for needles in a haystack. The unmethylated regions in a genome are highly stable during vegetative development and can reveal the locations of potentially expressed genes or cis-regulatory elements. This approach provides a framework towards complete annotation of genes and discovery of cis-regulatory elements using methylation profiles from only a single tissue.

## Introduction

There is a rapidly growing knowledge of the genome structure and sequence for many organisms. However, to fully utilize this resource, it is critical to identify and annotate the functional elements within the genome. In particular, there are two major challenges in providing high quality annotations of functional elements in complex eukaryotic genomes; correctly identifying functional genes and identification of cis-regulatory elements (CREs).

The challenge of identifying functional genes harkens to the oft-asked oral preliminary thesis exam question - what is a gene? This question seems to lack a good singular answer that can be applicable across different eras of genetics. While classical definitions of a gene were commonly based on mutant phenotypes it is clear that genetic redundancy or environment-specific phenotypic manifestations complicate our ability to identify phenotypes, even for functionally important genes. Genomics-based efforts to define gene-models are often based on a combination of evidence of transcripts and/or *ab initio* predictions. Yet, gene-models are best considered a hypothesis as to the existence of a gene (1). By and large, the majority of functional gene products can likely be captured based on identification of putative genes that are conserved in similar order among related species, often termed syntenic genes (1). However, there are also cases of functional genes that are created following gene duplication and/or transposition that are common in many plant genomes. One potential solution is to identify putative genes through genome-wide annotation and then to use chromatin features to filter the genes to highlight models that are more likely to retain function. These approaches have been applied in sorghum (2) and maize (3).

The problem of identifying potential CREs is even more challenging. In plants with large genomes, CREs can occur 10s-100s of kilobase pairs (kb) from their target genes (4). These regulatory regions, including gene-distal (hereafter, distal) CREs, have established roles in domestication and agronomic traits, for instance in *Zea mays* (maize) (5–11). Although only a handful of distal CREs have been characterized, recent studies suggest their prevalence in plants (4, 12–17). Yet, these regions do not necessarily produce easily detectable products (like transcripts) or have sequence features that can be identified such as protein-coding potential. Several approaches that survey accessible chromatin (12, 13, 17) or interactions of intergenic regions with gene promoters (18–20) are providing major insights for the identification of putative CREs. However, many of these technologies are specific to the tissue or cell type that is assayed. A complete understanding of the potential CREs within a particular species would require profiling of chromatin accessibility and/or chromatin interactions in a wide variety of tissues, cell types and conditions.

Although chromatin accessibility, histone modifications and chromatin interactions often show substantial variation in different tissues (12, 14, 21) the majority of DNA methylation patterns are quite stable in plant species especially during vegetative development (22–24) and in the face of environmental stress (25–28). There are well-characterized examples of specific changes in DNA methylation in endosperm tissues (29, 30) as well as in specific cell types in plant gametophytes (31–35). However, the majority of DNA methylation patterns are quite stable in different vegetative tissues, especially for DNA methylation in the CG and CHG contexts. In contrast, several studies have provided evidence for developmental or tissue-specific changes in CHH methylation (36–41). It should be noted however that the majority of these examples point to cases in which the level of CHH methylation at specific regions changes, but these regions often have some level of CHH methylation in all tissues.

Prior studies have found that the majority of regions of chromatin accessibility are hypomethylated (4, 12, 13, 17). Here, we reverse the approach and identify the unmethylated regions (UMRs) of the maize, barley, sorghum, rice and brachypodium genomes and compare the genomic distribution of UMRs with tissue-specific chromatin accessibility and provide evidence for functional roles of UMRs. We demonstrate that unmethylated regions of the genome, particularly in plant species with large genomes, provide useful information for identification of functional genes and CREs. This improves annotation of complex crop genomes and provides clear hypotheses about the portions of the genome that likely contain functional elements.

## Results

In general, the maize genome is highly methylated with only a small portion of the genome lacking DNA methylation (42–44). Deep whole genome bisulfite sequencing (WGBS) was performed on seedling leaf tissue of the inbred B73, which generated ∼930 million reads providing approximately 28x projected raw average coverage per strand (15.7x per cytosine average coverage following alignment and quality filtering) (Dataset S1). DNA methylation levels in the CG, CHG and CHH context were determined for each 100bp tile of the maize genome. While some regions lack cytosines in this context or could not be assessed due to lack of uniquely mapping reads, we were able to obtain DNA methylation estimates for 16.03 million tiles - ∼1.6 Gb - that contained at least two cytosines and an average of at least 5X coverage per cytosine per strand, representing 76.1% of the maize genome. A visual examination of a representative ∼100kb region containing two genes revealed that the majority of tiles are highly methylated; however, there are examples of unmethylated regions near syntenic genes and in distal regions (Figure 1A). Across the maize genome, 8.19% of the 100bp tiles with data - 131 Mb - had very low (<10%) or no detectable DNA methylation in any sequence context, termed unmethylated tiles (UMTs) (Figure S1A). We developed a framework to identify the unmethylated regions (UMRs) in a genome by first hierarchically categorising each tile into one of six methylation domains (see methods and Dataset S2), then merging adjacent unmethylated tiles into unmethylated regions. We restricted our analysis to 107,583 UMRs of at least 300bp, accounting for 5.8% of the maize genome (Figure S1A, Dataset S3b). These UMRs include many examples within genes, in gene proximal (within 2kb) regions and in distal regions at least 2kb from the nearest gene (Figure 1A-B). A more detailed investigation of the types of features overlapping with UMRs revealed significant enrichment for syntenic genes and depletion within intergenic regions (Figure S1C). Only a small proportion (10.2%) of the unmethylated 100bp tiles are found to overlap a variety of transposable elements (TE) from different orders (Figure S1D), but given the expectation that TEs are highly methylated it was interesting to note that there are 20,232 UMRs in total (18.8%) that overlap maize TEs.

**Fig 1.**
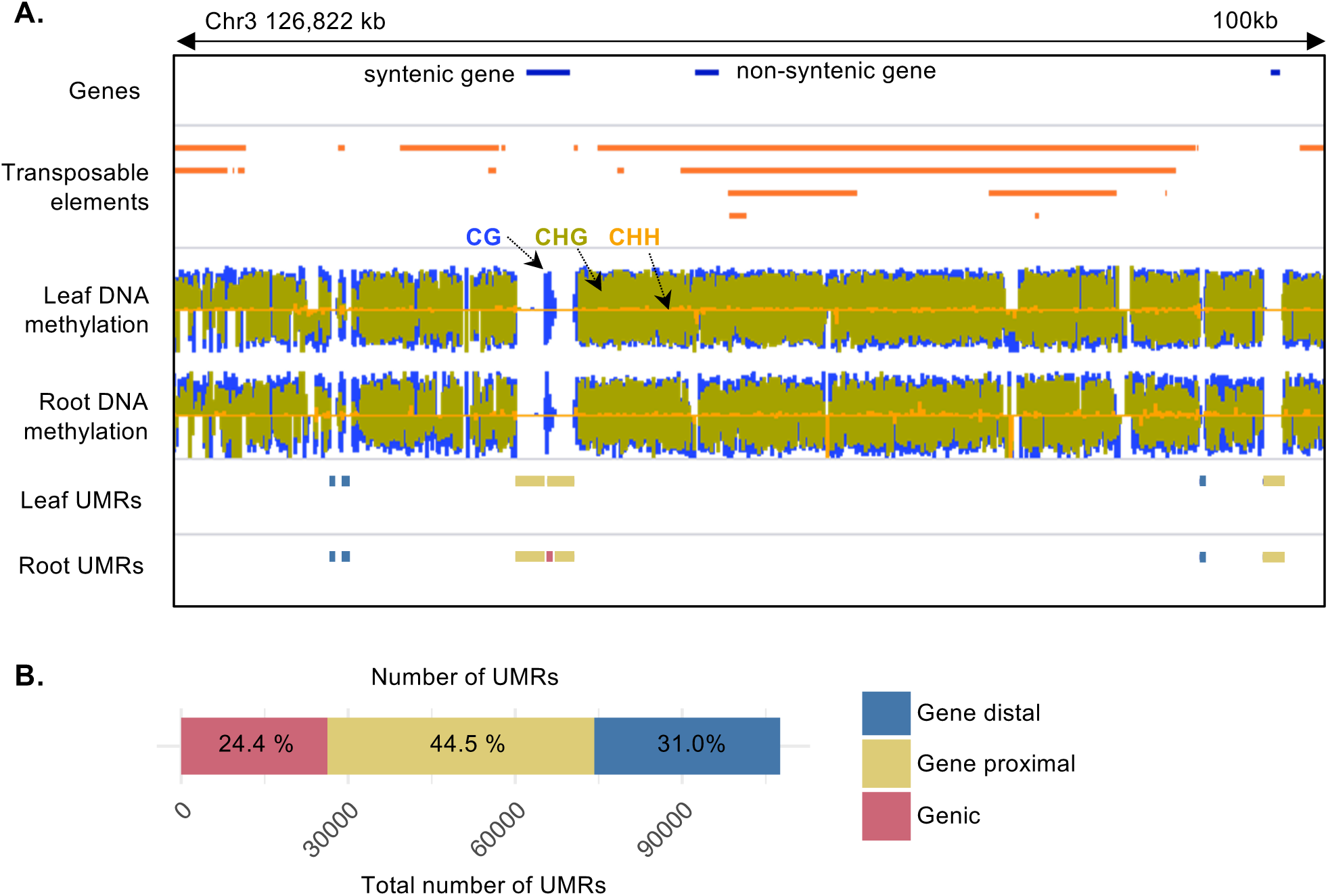
Identifying unmethylated regions in the maize genome. **A**. Example distribution of unmethylated regions (UMRs) in a 100 kb locus of the maize genome. The syntenic gene is hsp70-4 (Zm00001d041550). **B**. Genomic distribution of UMR. Proximal UMRs defined as those that overlap a 2kb window upstream of the TSS or 2kb downstream of the TTS (44.5%, 47,910), genic are entirely within the gene locus boundaries (24.4%, 26,298) and distal are >2kb from a gene (31.0%, 33,375). DNA methylation tracks for each context are indicated by the arrows colored as follows: blue CG; green CHG; orange CHH. Where UMRs overlap multiple genomic features, the location is annotated by hieratically categorization: proximal > distal > genic.

### Comparisons of unmethylated and accessible portions of the maize genome

There is evidence for enrichment of functional elements within accessible chromatin regions (ACRs) in maize (12, 13, 17). Several studies have found that these accessible regions tend to be hypomethylated (12, 13, 17). A comparison of the UMRs that are identified in B73 seedling leaf and root tissue using available WGBS data (Dataset S1)(45), reveals very few changes in UMRs between independent leaf samples or between tissues (Figure 1A and Figure 2A). There are ∼3-4% of UMRs identified solely in one of the samples; however, the vast majority of these were due to missing data in the other sample (Figure 2B). Less than 0.1% of the UMRs from one tissue are classified as methylated (Figure S1) in the other tissue, suggesting relatively infrequent changes in the UMRs among vegetative tissues in maize. Prior studies have found very few examples of major changes in CG or CHG methylation among vegetative tissues in maize or other plants (23, 24, 43). In contrast, chromatin accessibility profiles (Dataset S4) from three distinct tissues show significant variability (Figure 2C).

**Fig 2.**
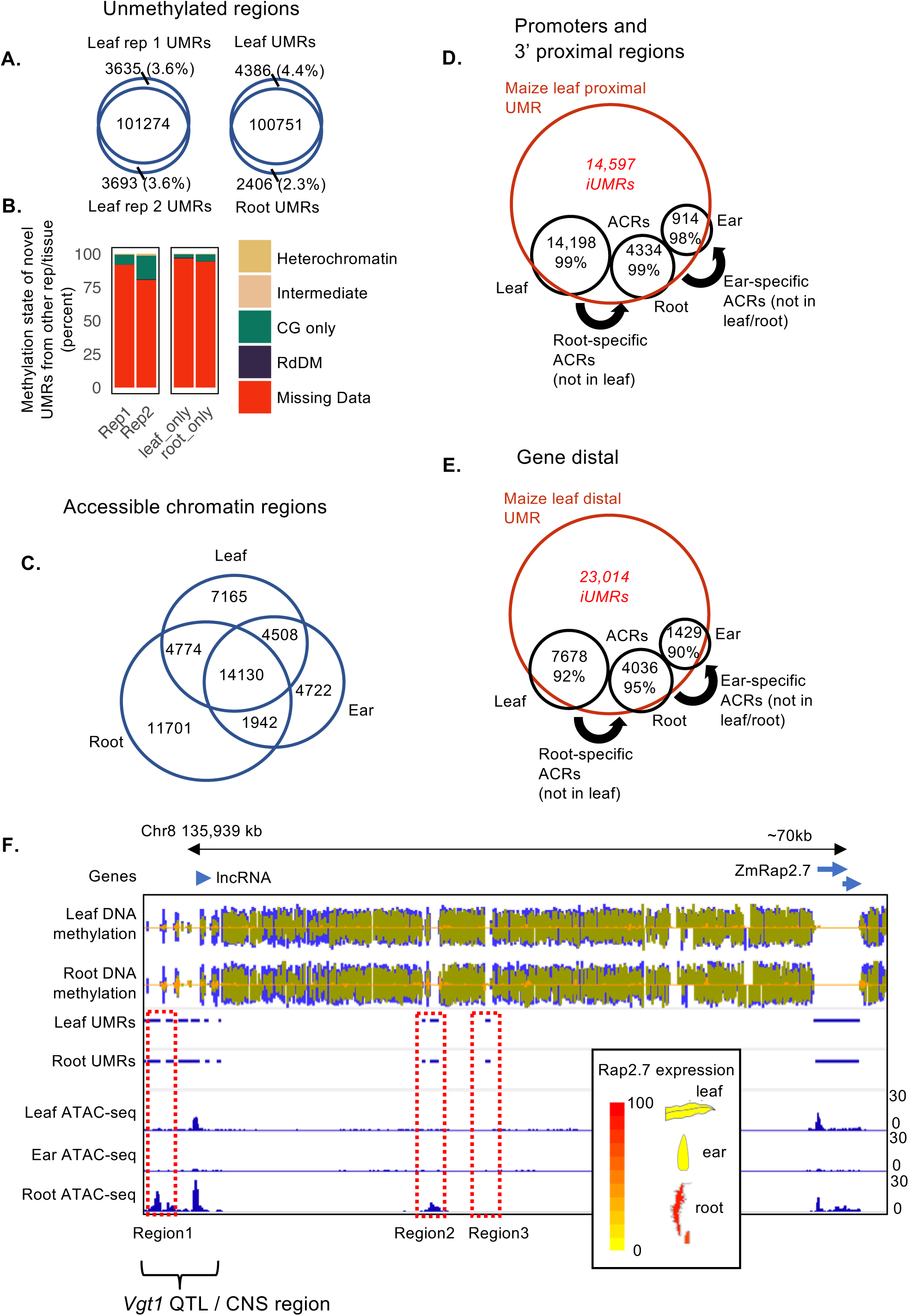
Comparisons of unmethylated and accessible portions of the maize genome. **A**. Overlap of unmethylated regions (UMRs) in two independent maize seedling leaf samples, “Rep 1” and “Rep 2”, and a seedling root sample. The percent of UMRs uniquely identified in one of the samples is listed in parentheses. **B**. For UMRs uniquely identified in one of the samples in A, the methylation data (methylation domain type) in the corresponding sample is displayed. For definitions of methylation domains see methods and Table S2. For example, over 80% of the UMRs uniquely identified in Rep2 have missing data in Rep1. **C**. Overlap of accessible chromatin regions (ACRs) from maize leaf (30,577), root (32,547) and ear (25,302). **D-E**. Gene proximal D., and gene distal E., UMRs from maize leaf capture the majority of both leaf ACRs as well as root- and ear-specific ACRs. Leaf UMRs were first overlapped with all leaf proximal, D; or distal, E; ACRs and the percent of ACRs overlapping is listed. Next, “root-specific” (not in leaf) ACRs were overlapped with leaf UMRs; then “ear-specific” (not in leaf or root) ACRs were overlapped with leaf UMRs. UMRs that do not overlap a leaf, root or ear ACR are inaccessible (iUMRs) and listed in red. **F**. Example leaf iUMRs that mark regions that become accessible in other tissues. Three **e**xample leaf iUMRs regions in the 70kb genomic region upstream of the rap2.7 transcription factor. The known vgt1 cis-regulatory region is marked by parenthesis. Region 1 and region 2 are examples of leaf iUMRs that are inaccessible in leaf but accessible in root tissue. Scale on the ATAC-seq tracks represent total read counts. The inset displays relative gene expression FPKM of rap2.7 in leaf, root and ear from the maize eFP browser.

Given the different dynamics in tissue-specific chromatin accessibility and tissue-specific DNA methylation, we were interested in exploring if UMRs from a single tissue could capture and predict potential ACRs in multiple tissues or conditions. We assessed the overlap of the seedling leaf UMRs with ACRs from three different tissues, including leaf and ear ACRs identified by Ricci, Lu, Ji, et al (12) and ACRs identified in root (Dataset S3), for gene proximal (Figure 2D) and gene distal regions (Figure 2E). As expected, the vast majority of ACRs overlap with UMRs. Over 99% of the promoter ACRs and 92% of the distal ACRs identified in seedling leaf tissue overlap with a UMR defined in seedling leaf tissue. Interestingly, when we focus on ACRs that are found in root tissue (but not in leaf) or in ear tissue (but not in leaf/root) we find that the vast majority of these are unmethylated in leaf tissue as well, despite being inaccessible in leaf (Figure 2D-E). Examination of DNA methylation and ATAC-seq data for several UMRs that exhibit accessibility solely in non-leaf tissues supports the observation of tissue-specific ACRs that are stably unmethylated (Figure 2F, Figure S2 and Figure S3). In two cases of classic maize genes, *tb1* (8) and *ZmRap2.7* (5) with defined long-distance enhancers, we find that UMRs are stable in multiple tissues, including in leaf tissues, where these genes are not appreciably expressed. In contrast, ACRs at distal regulatory regions and gene proximal regions only occur in tissues with expression for both genes (Figure 2F, Figure S2 and Figure S3). Combined, these observations suggest that UMRs defined on a single tissue may capture regions with potential for accessibility in a variety of cell types or tissues, thus providing a prediction of putative functionality.

### UMRs are indicative of expression potential of genes

To investigate accessibility dynamics of UMRs, UMRs defined on seedling leaf tissues were classified into two groups, accessible UMRs (aUMRs) and inaccessible UMRs (iUMRs), depending on the chromatin accessibility in seedling leaf tissue. In assessing the functional relevance of the aUMRs and iUMRs, we first focused on the UMRs found near gene transcription start sites (TSSs). There are 32,196 UMRs that overlap with the proximal region of maize genes (within 2kb upstream or 1kb downstream of the TSS) and 12,867 ACRs within these regions. Nearly all (>98%) of these ACRs overlap with an UMR (Figure 3A). However, 60.7% of UMRs that are located near gene TSS do not overlap an ACR. Considering recent work that demonstrated DNA methylation levels near the ends of the genes could predict expressibility of genes (3), we hypothesized that genes that are actively expressed in the tissue used for documenting accessibility (seedling leaf) would be enriched for ACRs/aUMRs, while genes expressed in other tissues and silenced in seedling leaf would be enriched for iUMRs. To do so, we gathered B73 RNA-seq data across more than 240 different samples from tissues, conditions and developmental stages; including seedling leaf RNA-seq data generated by Ricci, Lu, Ji, et al (12) (Dataset S5). All maize genes that are located in syntenic positions relative to other grasses (46) were classified as “leaf expressed” if they were detected at >1 counts per million (CPM) in seedling leaf tissue. The remaining genes were classified as “other tissue” if they were detected in at least one of the other tissues (>1CPM) or classified as “not expressed”. We then examined the methylation and accessibility of the promoters (defined as region 2 kb upstream of TSS to 1 kb downstream of TSS) of these genes (Figure 3B). In some cases, the lack of properly annotated TSSs for some gene models will lead to potential issues as promoter proximal regions will not be accurately defined. Syntenic genes that are expressed in leaf are enriched for aUMRs in the promoter proximal region (Figure 3B). However, there are almost as many genes expressed in this tissue that contain an iUMR and these may reflect examples in which the ACR region was too small to be effectively detected using ATAC-seq or expressed with limited accessibility (Figure 3B). Very few genes with leaf expression lack UMRs and ACRs (Figure 3B). Syntenic genes that are expressed in other tissues are less likely to contain an aUMR but frequently contain iUMRs (Figure 3B). A total of 1,323 genes expressed in other tissues (21.8% of “other tissue expressed”) contained an aUMR in their promoters, representing genes that are possibly poised in leaf for expression, have unstable transcripts or high transcript turnover, or contain silencing *trans*-factors in their promoters precluding their activation. We also identified cases where genes expressed in other tissues have inaccessible, but unmethylated promoters in leaf tissue that become accessible in other tissues, such as *NAC-transcription factor 114* (*nactf114/cuc3*) (Figure 3C). This gene is silent in leaf tissue but expressed in ear tissue (Figure 3D), yet its promoter is already unmethylated in leaf. Genes that are never detected as expressed are much less likely to contain aUMRs or iUMRs and more likely to be non-syntenic (Figure S4). If we assess all genes with an aUMR, we find that the majority are expressed in seedling leaf tissue. In contrast, genes with iUMRs are constitutively expressed, and are depleted of universally silenced genes. Genes that lack both iUMRs and aUMRs in leaf tissue are very rarely expressed in any tissue (Figure 3B) and are highly enriched for non-syntenic genes (Figure S4).

**Fig 3.**
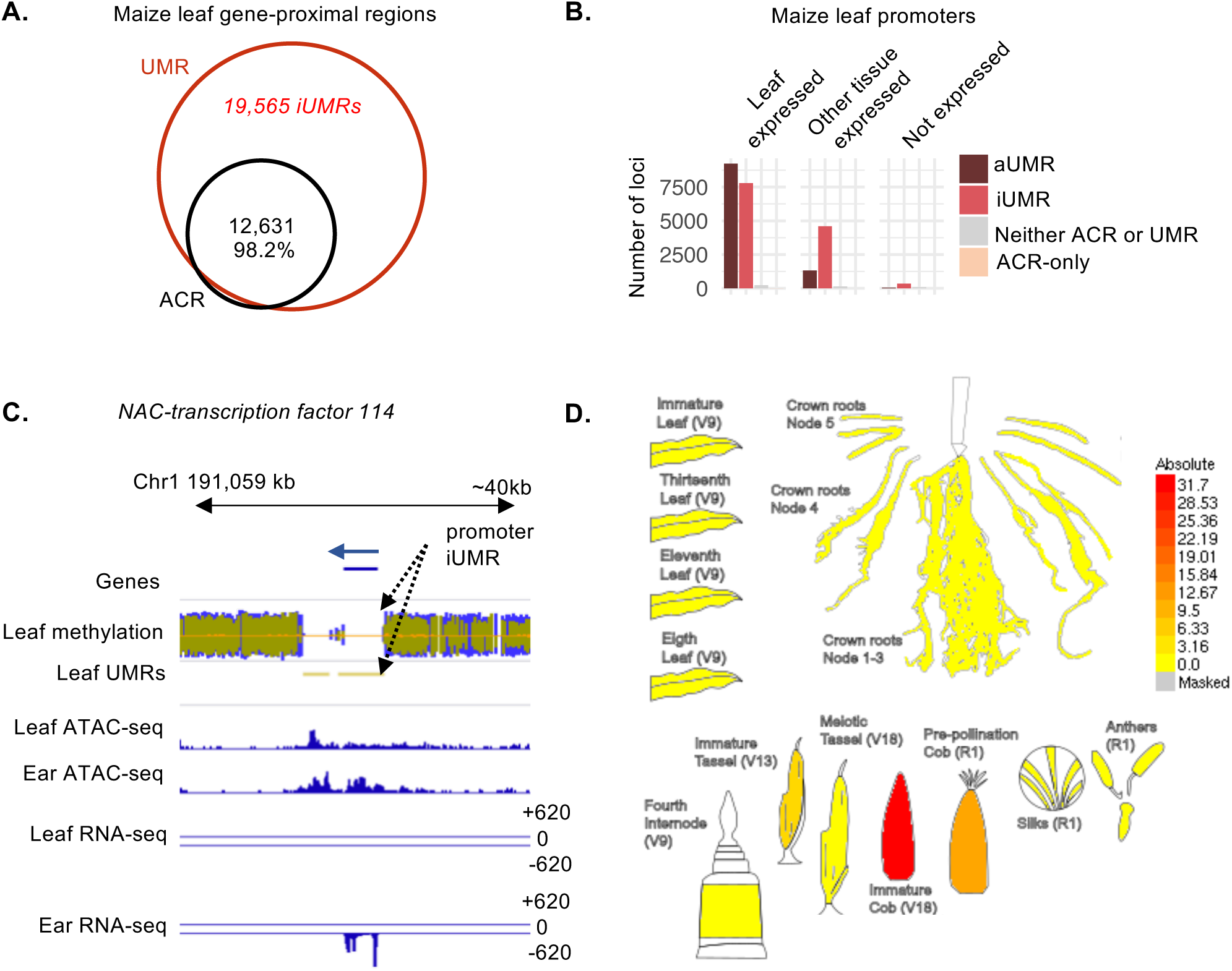
Gene proximal “promoter” unmethylated regions in maize. **A**. Unmethylated regions (UMRs) in gene promoters overlapped with accessible chromatin regions also found in gene promoter (ACRs). The percent of ACRs overlapping UMRs is 98.2%. Promoters or TSS proximal UMRs defined as those overlapping the region 2 kb upstream and 1 kb downstream of annotated TSS. **B**. Relationship between expression of a gene and promoter accessibility and methylation. Genes were defined as “leaf” expressed if expression was greater than 1 count per million (CPM) in leaf tissue; “other tissue” expressed if greater than 1 CPM in a tissue other than leaf but less than 1 CPM in leaf; or “not expressed” if expression less than 1 CPM in all tissues (only maize genes syntenic within the grasses shown). **C**. Example of a leaf promoter iUMR that may mark a gene for expression in another tissue. The promoter of the *NAC-transcription factor 114* (*nactf114*) Zm00001d031463 is unmethylated but inaccessible in leaf (iUMR black arrows). The promter region becomes accessible in ear and the gene is expressed in ear (RNA-seq). **D**. The relative expression (FPKM) profile of cuc3 in representative tissues from the maize eFP browser.

### Leaf UMRs are enriched for transcription factor binding sites

The concept that unmethylated regions from a single tissue can reflect sites with regulatory potential in diverse developmental stages or tissues suggests that UMRs from a single tissue could predict potential transcription factor (TF) binding sites, even for TFs only expressed in other tissues. We tested this concept in two different ways. First, we used the combined DAP-seq profiles for 32 maize TFs (12, 47) (Dataset S6). While these DAP-seq enriched regions only represent ∼1/10 of the genome, they account for 73% of the unmethylated regions and are enriched for both iUMRs and aUMRs (Figure 4A). Relative to a set of randomized control regions, there is evidence for enrichment (p < 0.001) of both aUMRs and iUMRs within the regions identified by DAP-seq (Figure 4A). Second, we used ChIP-seq data for five maize TFs, FASCIATED EAR4 (FEA4), KNOTTED1 (KN1), OPAQUE2 (O2), RAMOSA1 (RA1) and PERICARP COLOR1 (P1) (48–52). Notably, none of these TFs are highly expressed in seedling leaf tissues (Figure 4B and Figure S5-S10), but are expressed in other tissues or developmental stages. We were interested in assessing whether the binding sites for these TFs were unmethylated and inaccessible in the absence of their expression (note the leaf tissue used for methylation profiling comprises a 5cm section from the leaf, so excludes all tissue from the shoot apex including the meristem). For each TF, the number of ChIP-seq peaks that overlap with aUMRs and iUMRs is greater than expected by chance based on comparison to a set of randomly selected regions (Figure 4B, p < 0.001). However, the relative enrichment is quite variable, as some TFs such as FEA4 and KN1 show major enrichments but less enrichment for O2, RA1 and P1 (Figure 4B). This could reflect technical variation in the quality of the ChIP-seq datasets or may reflect differences in the potential for some TFs to bind methylated or unmethylated DNA. While some of the TF ChIP-seq peaks are found within aUMRs there are many that are found within UMRs that are inaccessible in leaf tissue. Indeed, relative to the expected number of iUMRs we find a significant enrichment (p < 0.001) for ChIP-seq binding peaks within UMRs for all of the 5 TFs. The observation that these binding sites are highly enriched for iUMRs suggests that the UMRs from this tissue can predict potential binding for these TFs in other developmental stages.

**Fig 4.**
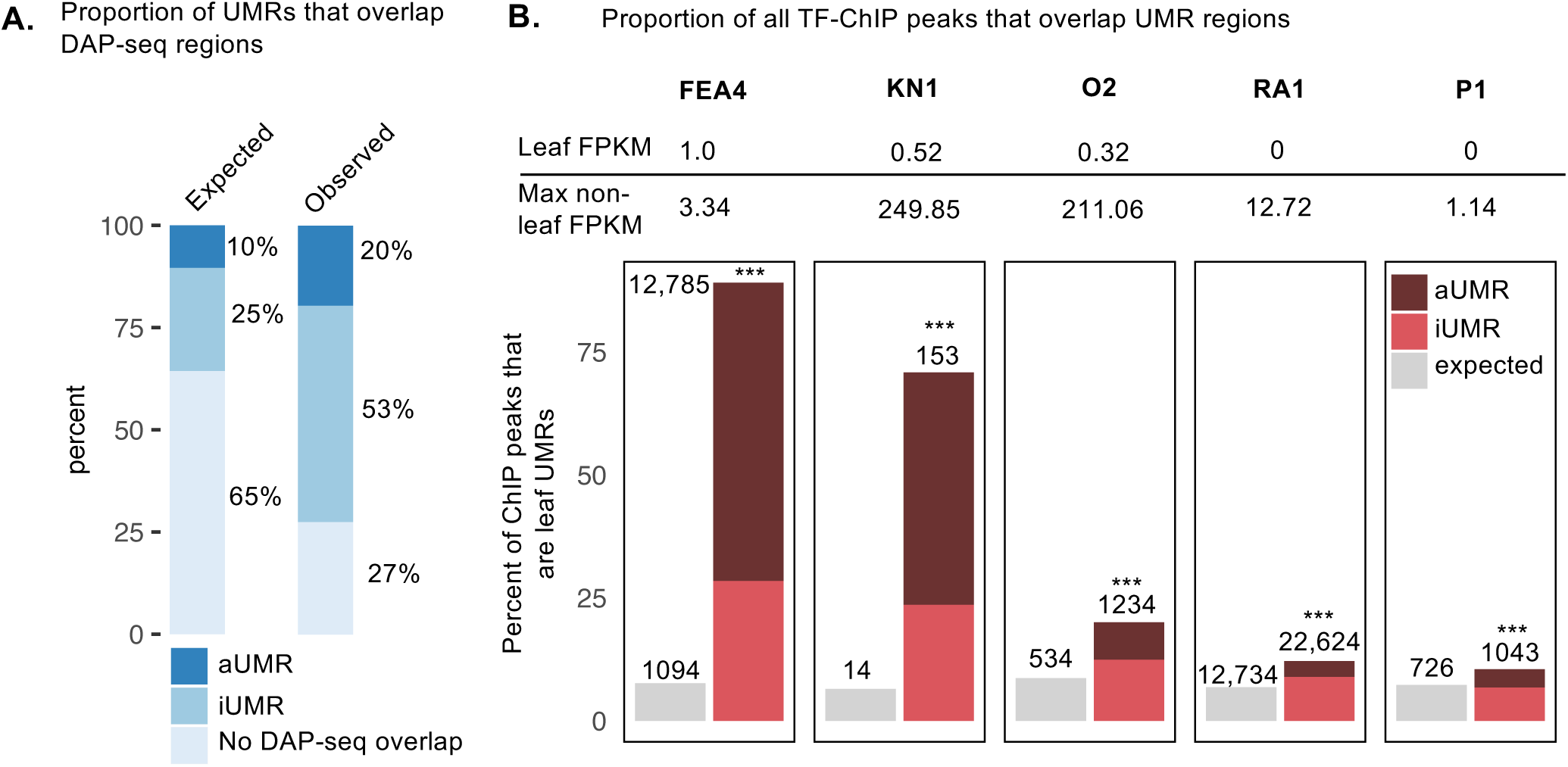
Transcription factor binding site enrichment in UMRs. **A** The expected and observed proportion of UMRs that overlap transcription factor (TF) binding sites identified using DAP-seq. UMRs that overlap TF binding sites are divided into accessible (aUMRs) and inaccessible (iUMRs) UMRs. **B**. Overlap of leaf UMRs with TFs binding sites determined using ChIP-seq for TFs which are expressed and function in non-leaf tissues. The expression of each TF in leaf and non-leaf tissues is listed about each plot. The tissues with maximum expression for each tissue is: *FEA4*, stem and SAM; *KN1*, immature cob; *O2*, endosperm; *RA1*, immature cob; *P1*, meiotic tassel. Expected ratios determined using random set of regions of equal number and size to the relevant contrast. Overlap *** = p < 0.001, based on 1000 permutations of randomised control regions.

### Evidence for enrichment of distinct chromatin features at distal UMRs

Cloning of agronomically important QTL has revealed several examples of important distal cis-regulatory regions that control expression of genes that are 10s-100s of kb away (5–8). Recent studies in maize have found evidence for many putative distal CREs based on accessible chromatin, chromatin modifications and 3-dimensional chromatin interactions (4, 12, 17–19). There are many distal aUMRs and iUMRs that are located at least 2kb from the nearest annotated gene (Figure 1B). These capture the majority of distal ACRs identified by ATAC-seq, even when these ACRs are not found in seedling leaf tissue (Figure 2E). We were interested in assessing whether the lack of DNA methylation at these distal regions was associated with unique chromatin profiles or function, especially for the iUMRs. The chromatin modifications from B73 seedling leaf tissue profiled by Ricci, Lu, Ji, et al (12) were used to compare the chromatin within and surrounding distal iUMRs and aUMRs with a set of random intergenic regions (Figure 5A). Analysis of ATAC-seq data from seedling leaf tissue confirmed the lack of accessible chromatin at iUMRs (Figure 5A). Both aUMRs and iUMRs exhibit altered profiles of many chromatin modifications relative to control regions both within the UMR and the flanking 1kb regions. The most striking difference between aUMRs and iUMRs is observed for H3K4me1; aUMRs tend to have quite low levels of this modification and are depleted for this mark in flanking regions. In contrast, iUMRs show a strong enrichment for H3K4me1 (Figure 5A). The majority of the other modifications examined exhibit similar trends for both iUMRs and aUMRs (Figure 5A). For some modifications, such as H3K4me3, H3K27ac, K3K9ac and H3K56ac, there are slightly stronger enrichments for the aUMRs (Figure 5A). In other cases, such as H3K27me3, the profiles are similar but the iUMRs have stronger enrichments. Both aUMRs and iUMRs are deleted for H3K9me2 but the depletion is stronger within the UMR relative to flanking regions of aUMRs (Figure 5A).

**Fig 5.**
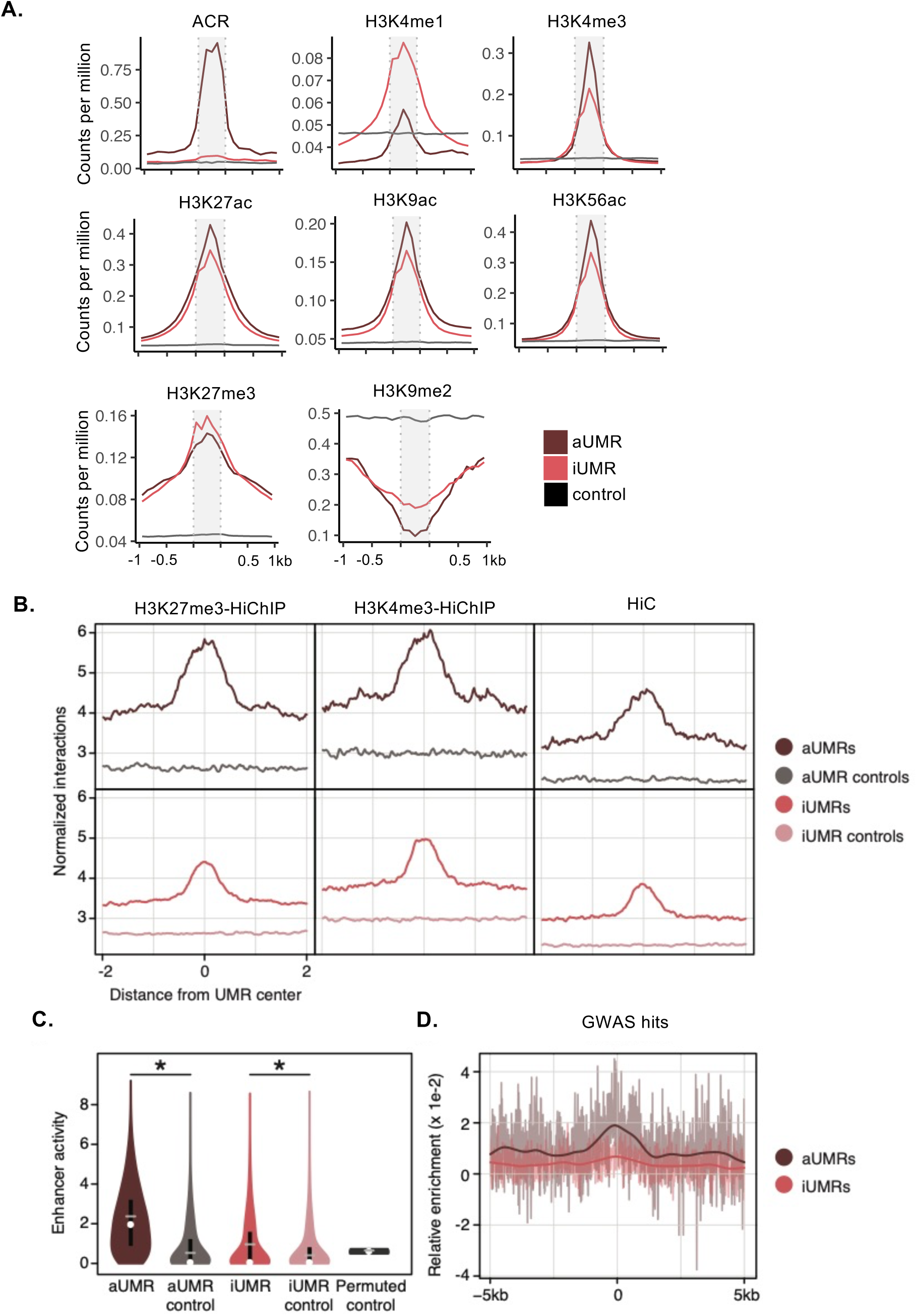
The chromatin profile of gene-distal iUMRs. **A**. The average enrichment of chromatin modifications over aUMR and iUMRs. Noramlised read abundance in counts per million for ATAC-seq (ACRs) and ChIP-seq for histone modifications is averaged over UMRs and the 1kb upstream and downstream. The UMR region is indicated by the shaded grey box. Standard error is overlayed in a lighter shade. **B**. Metaplot averages of normalized interaction tags from H3K27me3 Hi-ChIP, H3K4me3 Hi-ChIP, and Hi-C in 4-kb windows centered on iUMRs, aUMRs, and their respective controls. **C**. Distribution of enhancer activities (log2[RNA fragments per million / input fragments per million]) for aUMRs, iUMRs, their respective control regions, and averages from 10,000 Monte Carlo permutations of random intergenic regions. *denotes P<5e-108. **D**. Metaplot of relative enrichment of significant GWAS hits in 10-kb windows centered on iUMRs and aUMRs.

We proceeded to use several metrics developed by Ricci, Lu, Ji, et al (12) to investigate chromatin interactions and potential enhancer function for the iUMRs and aUMRs. We compared the proportion of iUMRs and aUMRs that overlap with HiC, H3K4me3-HiChIP or H3K27me3-HiChIP loop edges (Figure 5B, Figure S11). In each case we compared these to an associated control set of randomized intergenic regions. The iUMRs show nearly the same level of enrichment as the aUMRs suggesting that these regions are frequently making contacts with other regions and participate in chromatin looping. STARR-seq assays were performed by Ricci, Lu, Ji, et al (12) to assess the potential for ACRs to provide functional enhancer activity in maize leaf protoplasts. Given that many of the iUMRs are associated with genes that are expressed in other tissues or ChIP-seq peaks for TFs that are not expressed in leaf tissue, we did not expect the same level of enrichment for enhancer activity in protoplasts from leaf tissue. While the iUMRs show substantially less enhancer activity based on leaf protoplast STARR-seq assays compared to aUMRs we do still see significant enrichment relative to control regions (Wilcoxon rank sum test, p-value < 3.7e-108; and 10,000 permutations, empirical p-value < 1e-4) of matched intergenic sites (Figure 5C). We also assessed the frequency of GWAS hits at the iUMRs relative to aUMRs (Figure 5D). While the aUMRs show significant increase for GWAS hits there is less enrichment for iUMRs. Overall, these analyses suggest that iUMRs and aUMRs have unique chromatin profiles relative to the other distal intergenic regions and that the iUMRs often participate in chromatin loops. However, these iUMRs do not show the same level of enrichment for GWAS hits as aUMRs, which may reflect a greater level of conservation or signal loss due to inclusion of non-functional sites.

### The utility of UMRs for annotation and discovery in large genomes

These analyses were initially focused on maize given the availability of other datasets that could be used to assess potential functions and roles of iUMRs. However, we predict that similar numbers of iUMRs and aUMRs would be identified in other cereals and grasses. We gathered DNA methylation and chromatin accessibility data for four other grasses (Dataset S1); barley (*Hordeum vulgare*) (53), sorghum (*Sorghum bicolor*) and brachypodium (*Brachypodium distachyon*) (54), and rice (*Oryza sativa*) (55). These species vary substantially in genome size with some species <500Mb and others >4GB (Figure 6A). The genome size that could be assessed for DNA methylation varied with >1.5GB for maize and barley and <400Mb for rice and brachypodium. Despite these major differences in total genome size and the size of the genome for which DNA methylation could be profiled, we find roughly similar amounts of UMRs across all profiled species (Figure 6A, Dataset S7 - Dataset S10). This suggests that the total genome space of UMRs is relatively constant despite dramatic changes in overall genome size. This is consistent with the finding that the total accessible space (ACRs) is similar in genomes of different sizes (4). The distribution of genic, proximal and distal UMRs varies between species but is related to genome size (Figure 6B). The large genomes have more examples of distal UMRs and relatively fewer proximal UMRs compared to small genomes. This likely reflects the higher gene density in smaller genomes and reduction in the amount of genome classified as distal intergenic. If we assess the proportion of all genic, proximal and distal space in each genome that is classified as UMR, we find that the amount of genic space within UMRs is quite similar for all species (Figure 6C). In contrast, the proportion of distal space that is classified as UMR is much lower in species with large genomes (Figure 6C). This highlights the potential of DNA methylation to reveal the subset of potentially functional intergenic space, particularly in large genomes.

**Fig 6.**
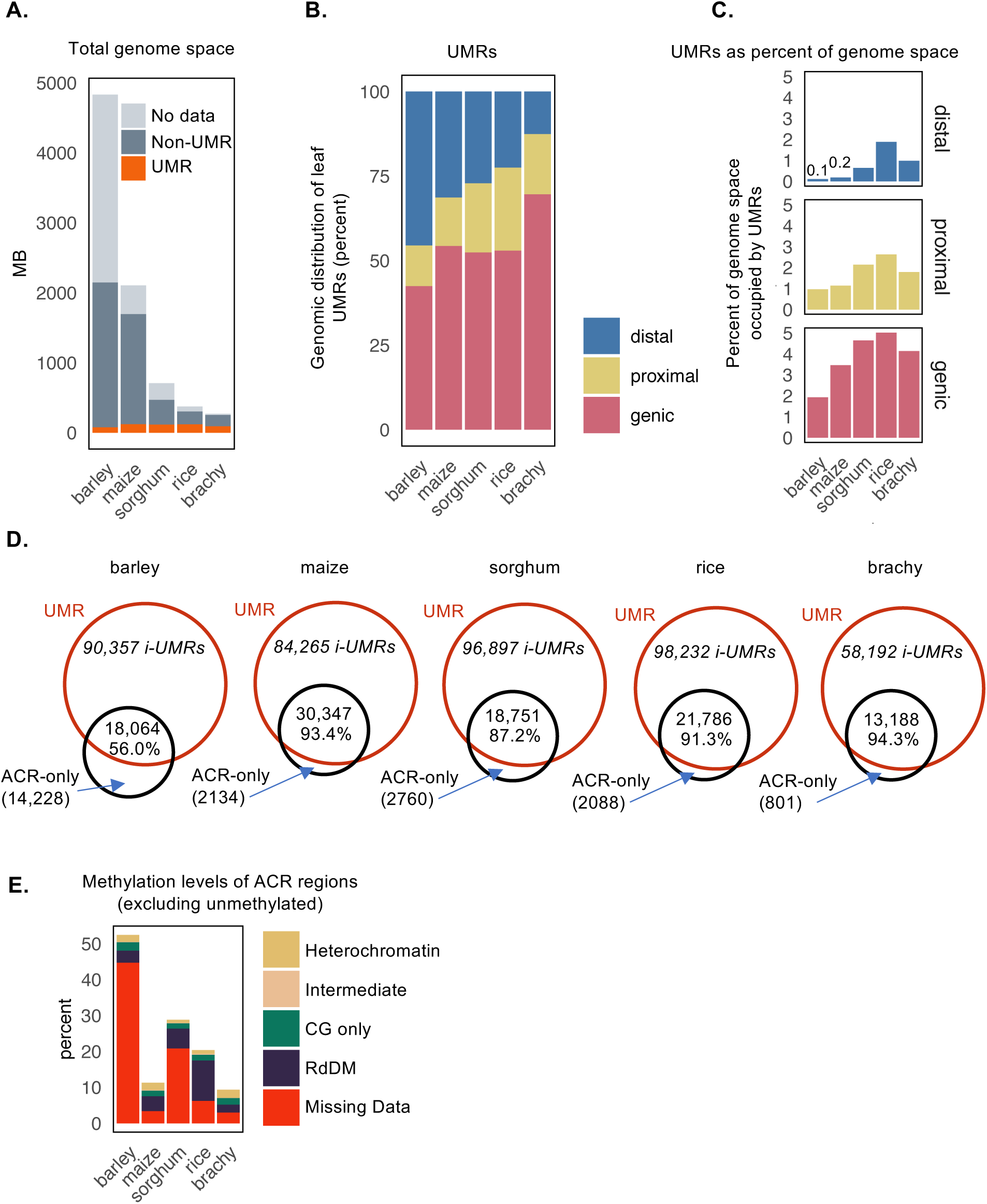
Similarity between the absolutes size and features of the unmethylated portion of cereal genomes. **A**. Comparative UMR analysis of 5 cereal genomes. The height of each bar represents the total size (MB) of each assembled genome, which is divided into the proportion of UMRs, non-UMR and regions lacking data. The total MB of UMRs in each genome is very similar. **B**. The genomic location of UMRs. Proximal overlap a 2kb window upstream of the TSS or 2kb downstream of the TTS, genic are entirely within the gene locus boundaries and distal are >2kb from a gene. **C**. The proportion of genic, proximal and distal regions of each genome comprised of UMRs. Larger genomes, such as maize and barely, have a much larger intergenic space and hence intergenic/distal UMRs are much smaller fraction. **D**. Overlap between UMRs and ACRs in each species, percentages refer to the percent of ACR that are unmethylated and captured by UMR profiling (not corrected for missing data). **E**. The methylation profile (distribution of methylation domain types) of ACR regions in each species, excluding unmethylated regions. For definitions of methylation domains see methods and Table S2. For example, over 43.6 % of the ACRs in barley overlap regions with missing methylation data, explaining the relative low overlap in D.

A comparison of the UMRs and ACRs for each species revealed substantial overlap, the same as was observed in maize (Figure 6D). In most species, the vast majority of ACRs occur within UMRs. The proportion of overlap is the smallest for barley. However, when we assessed ACRs failing to overlap with UMRs, we found that for all species the vast majority of these represent either unmethylated tiles that did not meet the criteria for UMRs or had missing data (Figure 6E). There are very few examples of ACRs in any of the species that are classified as having high levels of heterochromatic DNA methylation. The observation that some ACRs are not captured within the classified UMRs due to missing data for DNA methylation highlights the importance of deep coverage methylation datasets for use in annotation of UMRs. The barley methylome dataset is only ∼4.6X (Dataset S1) and this results in a substantial amount of genomic space that does not have sufficient sequencing depth for accurate classification of the DNA methylation state.

## Discussion

The annotation of genomes remains a difficult problem, especially the discovery of putative regulatory elements. Documenting tissue- or cell-specific expression levels and chromatin states has been successfully applied to improve annotations using ENCODE-like approaches (56–59). However, generating comprehensive atlases of expression or chromatin in many cell types and conditions can be experimentally challenging and costly. Here we suggest that identification of the unmethylated portions of crop genomes from a single tissue can help provide fairly complete catalogues of potential regulatory elements and expressed genes across many developmental stages. The advantage to this approach is that it can be performed on a single, easily harvested tissue type. As we noted in our comparison of species, it is important to generate a relatively deep coverage dataset of DNA methylation to maximize the amount of the genome that is confidently classified as methylated or unmethylated. Even with deep coverage, monitoring both UMRs and ACRs within highly repetitive regions remains challenging. In maize, we are only able to profile methylation levels for ∼70% of the maize genome with a relatively deep coverage dataset. The remainder is too repetitive to allow unique mapping using short reads.

It is worth noting that this conceptual framework - using UMRs from a single tissue to develop a catalogue of potential regulatory elements and expressed genes - relies upon the stability of CG and CHG methylation in different cell types. While CG methylation can be quite variable in different cell types for mammals (60), the CG and CHG methylation patterns are dramatically more stable in plants. There are examples of dynamic CHH methylation in different tissues in plants (36–41); and there is also evidence that some specific cell types undergo substantial changes in the methylation during reproduction (31–33). However, the bulk of the genome exhibits an often underappreciated consistency in the patterns of DNA methylation among different vegetative tissues (22–24).

One application of the UMR framework is to identify genes that have potential for expression. In maize, and other crop genomes, it is difficult to discriminate transposon-derived gene fragments from true genes (1, 61). Annotation of genes requires a balancing between quality and comprehensiveness (61). In many cases the desire to have a relatively complete set of putative genes results in numerous pseudogenes being included in gene annotations. Prior work has shown that applying machine learning to the patterns of context specific levels of DNA methylation can classify genes with potential for expression (3). Here we show that the majority of genes that are detected as expressed (in a panel of >200 samples) contain an unmethylated region close to their annotated TSS. The presence of a UMR within the promoter of a putative gene can be used to indicate the potential for expression of the gene. Several other studies have implemented different approaches to use DNA methylation data to augment gene annotations (2, 3).

UMRs can also be used to discover potential regulatory elements. UMRs are enriched for TF binding sites based on DAP-seq or ChIP-seq datasets. It is especially noteworthy that this enrichment for TF binding is observed even though the UMRs are defined in a tissue-type where most of these TFs are very low expressed or silent. There are also many examples of tissue-specific ACRs that are equally unmethylated across multiple tissues. This suggests that in plant genomes the majority of regions with potential to be TF binding sites in some tissue, developmental stage or environment, are stably unmethylated. While the application of chromatin accessibility assays or TF-binding assays in a specific tissue can provide a very high quality representation of the active regulatory elements in that tissue; here, we show that it is possible to rapidly develop a far more complete set of potential regulatory elements through the analysis DNA methylation profiles from a single tissue.

The utility of a methylation filter to focus on unmethylated regions may be variable across different species. In species with relatively small genomes, for example < 500 Mb, and limited intergenic space the filtering power of focusing on unmethylated regions is likely diminished. In a species like *Arabidopsis thaliana* most genes are arranged in close proximity to other genes and the amount of the genome that could be masked as methylated is relatively small (62). In contrast, in species such as barley or maize with large genomes and low gene density the ability to focus on the unmethylated portions of the genome provides a powerful framework to distil an enormous genome down to a relatively small fraction of genomic space, highly enriched regions valuable for regulation or manipulation of plant traits.

## Methods

### WGBS data

Whole genome bisulfite sequencing samples are listed in Dataset S1. For deep coverage maize leaf data generated in this study, DNA was extracted from leaves of two-week-old V2 glasshouse grown maize B73 plants using the DNeasy Plant Mini kit (Qiagen). Six biological replicates were sampled for sequencing and later combined into a single data set. 1ug of DNA in 50ug of water was sheared using an Ultrasonicator (Covaris) to approximately 200-350 bp fragments. 20ul of sheared DNA was then bisulfite converted using the EZ DNA Methylation-Lightning Kit (Zymo Research) as per the manufacturer’s instructions and eluted in a final volume of 15ul. Then 7.5ul of the fragmented, bisulfite-converted sample was used as input for library preparation using the ACCEL-NGS Methyl-Seq DNA Library Kit (SWIFT Biosciences). Library preparation was performed as per the manufacturer’s instructions except each reaction was scaled by a half. The indexing PCR was performed for 5 cycles. Libraries were then pooled and sequenced on a HiSeq 2500 in high output mode 125bp paired end reads over multiple lanes at the University of Minnesota Genomics Centre. WGBS data generated in this study is available under accession GSE150929. Additional maize seedling leaf (SRR8740851) and seedling root (SRR8740850) samples were described previously (45) and downloaded from SRA PRJNA527657. WGBS data for other species was downloaded from SRA including barley leaf (*H. vulgare* accession Morex) SRR5124893 (53), sorghum leaf (*S. bicolor*, accession BTx623) SRR3286309 (54), rice leaf (*O. sativa*, accession Nipponbare) SRX205364 (55) and brachypodium leaf (*B. distachyon*, accession Bd21) SRX1656912 (54).

Sequencing reads were trimmed and quality checked using *Trim galore!* version 0.4.3, powered by *cutadapt v1.8.1* (63) *and fastqc v0.11.5*. For swift libraries, 20bp were trimmed from the 5’ ends both R1 and R2 reads as per the manufacturer’s suggestion. Reads were aligned with *bsmap v2.74* (64) *to the respective genomes with the following parameters -v 5* to allow 5 mismatches, *-r 0* to report only unique mapping pairs, *-p 1, -q 20* to allow quality trimming to q20; *Z. mays* genome and gene annotation AGPv4 (65) downloaded from MaizeGDB gramene version 36, *H. vulgare L*. from Ensemble (v.42), *S. bicolor v3.1.1* from JGI (phytozome v.12), *O. sativ*a v7.0 from (phytozome v.11), *B. distachyon* v3.1 (phytozome v.12). For barley, as per Beier et al. (66), to accommodate limitations of the Sequence/Alignment Map format split pseudomolecules with a size below 512 Mb were used (downloaded from DOI:10.5447/ipk/2016/36). All genomes used for WGBS analysis were identical to assemblies used for ATAC-seq and all other genomic analyses. Output SAM files were parsed with *SAMtools (67) fixsam*, sorted and indexed. *Picard MarkDuplicates* was used to remove duplicates, *BamTools filter* to remove improperly paired reads and *bamUtils clipOverlap* to trim overlapping reads so as to only count cytosines once per sequenced molecule in a pair for PE reads. The *methylratio.py* script from *bsmap v2.74* was used to extract per site methylation data summaries for each context (CH/CHG/CHH) and reads were summarised into non-overlapping 100bp windows tiling the genome. *bedtools* and the script *bedGraphToBigWig* (68) were used to prepare files for viewing in IGV (69). WGBS pipelines are available on github (https://github.com/pedrocrisp/springerlab_methylation).

### Identification of unmethylated regions

To identify unmethylated regions, each 100bp tile of the genome was classified into one of six domains or types, including “missing data” (including “no data” and “no sites”), “RdDM”, “Heterochromatin”, “CG only”, “Unmethylated” or “intermediate”, according to the hierarchy in Dataset S2 (also see Figure S1). Briefly, tiles were classified as “missing data” if tiles had less than 2 cytosines in the relevant context or if there was less than 5x coverage for maize or less than 3x coverage when comparing the different grass species (owing to lower coverage in some species); “RdDM” if CHH methylation was greater than 15%; “Heterochromatin” if CG and CHG methylation was 40% or greater; “CG only” if CG methylations was greater than 40%, “Unmethylated” if CG, CHG and CHH were less than 10% and “Intermediate” if methylation was 10% or greater but less than 40%. Following tile classification, adjacent unmethylated tiles (UMTs) were merged. To capture and combine any unmethylated regions that were fragmented by a short interval of missing data (low coverage or no sites), any merged UMT regions that were separated by “missing data” were also merged so long as the resulting merged region consisted of no more than 33% missing data. Regions less than 300 bp were removed (Figure S1) and the remaining regions defined as unmethylated regions (UMRs).

### ATAC-seq data

For maize root ATAC-seq data generated in this study, *Z. mays* (accession B73) were grown in soil for around 6 days at 25 °C under 16 h light–8 h dark. The staple roots were harvested and were used for experiments. ATAC-seq was performed as described previously (70). For each replicate, approximately 100 mg of maize staple roots were harvested and immediately chopped with a razor blade and placed in 2 ml of pre-chilled lysis buffer (15 mM Tris–HCl pH 7.5, 20 mM sodium chloride, 80 mM potassium chloride, 0.5 mM spermine, 5 mM 2-mercaptoethanol and 0.2% TritonX-100). The chopped slurry was filtered twice through miracloth. The crude nuclei were stained with 4,6-diamidino-2-phenylindole and loaded into a flow cytometer (Beckman Coulter MoFlo XDP). Nuclei were purified by flow sorting and washed in accordance with (70). The sorted nuclei (50,000 nuclei per reaction) were incubated with 2 μl of transposome in 40 μl of tagmentation buffer (10 mM TAPS–sodium hydroxide pH 8.0, 5 mM magnesium chloride) at 37 °C for 30 min without rotation. The integration products were purified using a NEB Monarch PCR Purification Kit and then amplified using Phusion DNA polymerase for 11 cycles (71). Amplified libraries were purified with AMPure beads to remove primers.

ATAC-seq raw reads were aligned as described before (4, 12). Raw reads were trimmed with *Trimmomatic v*.0.33 (72). Reads were trimmed for NexteraPE with a maximum of two seed mismatches, a palindrome clip threshold of 30 and a simple clip threshold of ten. Reads shorter than 30 bp were discarded. Trimmed reads were aligned to the *Z. mays* AGPv4 reference genome (65) using *Bowtie v.1.1.1* (73) with the following parameters: *‘bowtie -X 1000 -m 1 -v 2 --best –strata’*. Aligned reads were sorted using *SAMtools v.1.3.1* (67) and clonal duplicates were removed using *Picard version v.2.16.0* (http://broadinstitute.github.io/picard/). *MACS2* (74) was used to define ACRs with the *‘-keepdup all’* function and with ATAC-seq input samples (Tn5 transposition into naked gDNA) as a control. The ACRs identified by *MACS2* were further filtered using the following steps: (1) peaks were split into 50 bp windows with 25 bp steps; (2) the accessibility of each window was quantified by calculating and normalizing the Tn5 integration frequency in each window with the average integration frequency across the whole genome to generate an enrichment fold value; (3) windows with enrichment fold values passing a cutoff (30-fold) were merged together by allowing 150 bp gaps and (4) possible false positive regions were removed by filtering small regions with only one window for lengths >50 bp. The sites within ACRs with the highest Tn5 integration frequencies were defined as ACR ‘summits’. ATAC-seq data for maize leaf and ear were as described in Ricci, Lu, Ji, et al. (12) and the coordinates of accessible chromatin regions were downloaded from the GEO archive GSE120304. ATAC-seq data for barley (*H. vulgare* accession Morex), sorghum (*S. bicolor*, accession BTx623), rice (*O. sativa*, accession Nipponbare) and brachypodium (*B. distachyon*, accession Bd21) are as described in Lu et al. (4) and the coordinates of accessible chromatin regions were downloaded from the GEO archive GSE128434.

### Expression data

RNA-seq expression data for maize leaf and ear (12) as well as 247 samples for other maize tissues (75–84) were downloaded from NCBI Sequence Read Archive and processed as described in Zhou et al (85). Briefly, reads were trimmed by *Trim Galore!* and *Cutadapt* (63) and aligned to the B73 maize reference genome (AGPv4, Ensembl Plant release 32) using *Hisat2* (86). Uniquely aligned reads were then counted per feature by *featureCounts* (87). Raw read counts were then normalized by library size and corrected for library composition bias using the TMM normalization approach (88) to give CPMs (Counts Per Million reads) for each gene in each sample. Hierarchical clustering and principal component analysis clustering were used to explore sample cluster patterns. Pipeline scripts, normalization code, and expression matrices are available at Github (https://github.com/orionzhou/rnaseq/tree/master/data/11_qc/rnc01).

Synteny classifications (i.e., syntenic and non-syntenic) and assignment to maize sub-genomes were obtained from a previous study based on pairwise whole-genome alignment between maize and sorghum, downloaded from Figshare Schnable 2019: DOI:10.6084/m9.figshare.7926674.v1 (46). The eFP browser expression data was downloaded from bar (89) hosted on Maize GDB incorporating the maize expression datasets (82, 90). The expression of fea4, kn1, o2, ra1 and p1 in leaf was evaluated considering the samples: pooled leaves V1, topmost leaf V3, tip of stage 2 leaf V5, base of stage 2 leaf V5, tip of stage 2 leaf V7, base of stage 2 leaf V7, immature leaf V9, thirteenth leaf V9, eleventh leaf V9 eighth leaf V9, thirteenth leaf VT, thirteenth leaf R2.

### Transcription factor DAP-seq and ChIP-seq

DAP-seq profiles for 32 maize TFs (12, 47) (Dataset S6) were downloaded from the SRA as were ChIP-seq for 5 TFs: KNOTTED1 (KN1) (48), RAMOSA1 (RA1) (49), fasciated ear4 (FEA4) (50), Opaque2 (O2) (52), PERICARP COLOR 1 (p1) (51). Sequencing data were downloaded from NCBI using the SRA Toolkit and processed using the nf-core ChIP-seq pipeline (91). Briefly, reads were trimmed by *Trim Galore!* and *Cutadapt* (63) and aligned to the B73 maize reference genome (AGPv4, Ensembl Plant release 32) using *BWA* (92). *Picard* was used to merge alignments from multiple libraries of the same sample and then to mark PCR duplicates. Further filtering was employed to remove reads marked as duplicates, non-primary alignments, unmapped or mapped to multiple locations, having an insert size > 2kb or mapped with abnormal paired-end signatures (only one read out of a pair mapped, two reads mapped to two different chromosomes or in abnormal orientation, etc) using *SAMtools. MACS2* was used to call broad and narrow peaks (“*--gsize=2.1e9 --broad-cutoff=0.1*”) and calculate the FRiP scores (74). *HOMER* was used to annotate peaks relative to gene features. Genome-wide IP enrichment relative to control was calculated using *deepTools2* (93) and strand cross-correlation peak and ChIP-seq quality measures including NSC and RSC were calculated using *phantompeakqualtools* (94). Randomized control regions were generated using *bedtools shuffle* and comparison with UMRs evaluated with 1,000 permutations using *regioneR*. Pipeline scripts, QC files and peak calling results and annotation are available at Github (https://github.com/orionzhou/chipseq).

### Analysis of histone modifications

Chromatin immunoprecipitation followed by high-throughput sequencing (ChIP-seq) data for H3K3me1, H3K3me3, H3K27ac, H3K9ac, H3K56ac, H3K27me3, H3K9me2 chromatin modification reported by Ricci, Lu, Ji, et al (12) were downloaded from the GEO database accession GSE120304. Adapter sequences were removed from raw reads using *Trimmomatic version 0.33* with default settings. Quality filtered reads were aligned to the maize B73v4 genome using *bowtie 1.1.1* with the following parameters: *-m 1 -v 2 –best --strata --chunkmbs 1024 -S*. Only uniquely mapped reads were retained and duplicated reads were removed using *rmdup* module from *samtools version 0.1.19*. Output bam files were used to count the number of reads aligning to each 100bp window of the B73v4 genome. Counts were normalized per million mapped reads. Chromatin metaplots for gene distal aUMRs and iUMRs were generated using *bedtools closest* to associate each 100bp window with the nearest UMR, each window could only be associated with a single UMR. Average meta-profiles were calculated for 1kb flanking upstream and downstream of UMRs and relative distance was determined for the 100bp windows within the annotated UMR. A control set of UMRs was generated using *bedtools shuffle* randomizing the same length regions across the genome.

### Analysis of Hi-C, Hi-ChIP and STARR-seq data

Raw and processed Hi-C, Hi-ChIP and STARR-seq data were acquired from Ricci, Lu and Ji et al (12). Analysis of chromatin interaction frequencies overlapping UMRs was performed using *DeepTools* (93) using 4-kb windows centered on UMRs with 20 bp bins. Length and counted-matched intergenic control regions for iUMRs and UMRs were selected using bedtools shuffle, excluding UMRs, ACRs and B73 V4 annotated repetitive elements. Per base pair STARR-seq enhancer activities were estimated as the log2 ratio between RNA and input fragments scaled per million mapped fragments. UMR enhancer activities were then taken as the maximum signal from single-bp resolution STARR-seq tracks. Comparison of enhancer activities between UMRs and controls were evaluated with Wilcoxon rank sum test and verified using 10,000 permutations of matched intergenic regions. Empirical significance values from the Monte Carlo permutation were estimated as the number of independent permutations with mean enhancer activities greater than the mean of the cognate UMR distribution.

### GWAS data analysis

Genomic positions of significant GWAS hits from Wallace et al. (95) were converted to V4 coordinates using the *liftOver* bioconductor R package. GWAS hits and SNPs in the NAM founder lines (BAM files acquired from cyverse) were counted in 10kb windows centered on UMRs with 10bp bins using *DeepTools*. Genome-wide coverage of NAM founder lines was determined using *bedtools genomecov* and aggregated across founders. Bins below the 5% quantile of genome-wide coverage of the NAM founders were set to missing and excluded from calculation. Relative GWAS enrichment per bin was estimated as the ratio between the mean GWAS and SNP counts of UMRs subtracted by the ratio of the mean GWAS and SNP counts of control regions (Eq 1).

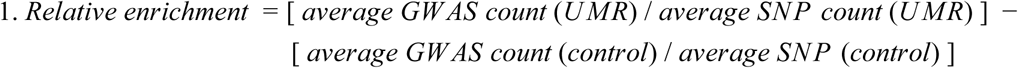

## Acknowledgements

This work was funded by grants NSF IOS-1934384 to NMS, NSF IOS-1802848 to NMS and NSF IOS-1856627 to RJS. PAC is the recipient of an Australian Research Council Discovery Early Career Award (project number DE200101748) funded by the Australian Government. JMN was supported by a Hatch grant from the Minnesota Agricultural Experiment Station (MIN 71-068). AM was supported by an NSF Postdoctoral Fellowship in Biology (NSF DBI-1905869). RS is a Pew Scholar in the Biomedical Sciences, supported by The Pew Charitable Trusts. The Minnesota Supercomputing Institute at the University of Minnesota provided computational resources that contributed to this research.

## Author Contributions

NMS, RJS and PAC conceived and planned the research. APM, JMN and ZL performed experiments. PAC, APM JMN, PZ and ZL conducted data analysis. PAC and NMS drafted the manuscript and all authors reviewed and commented on the manuscript.

---

**Fig S1.**
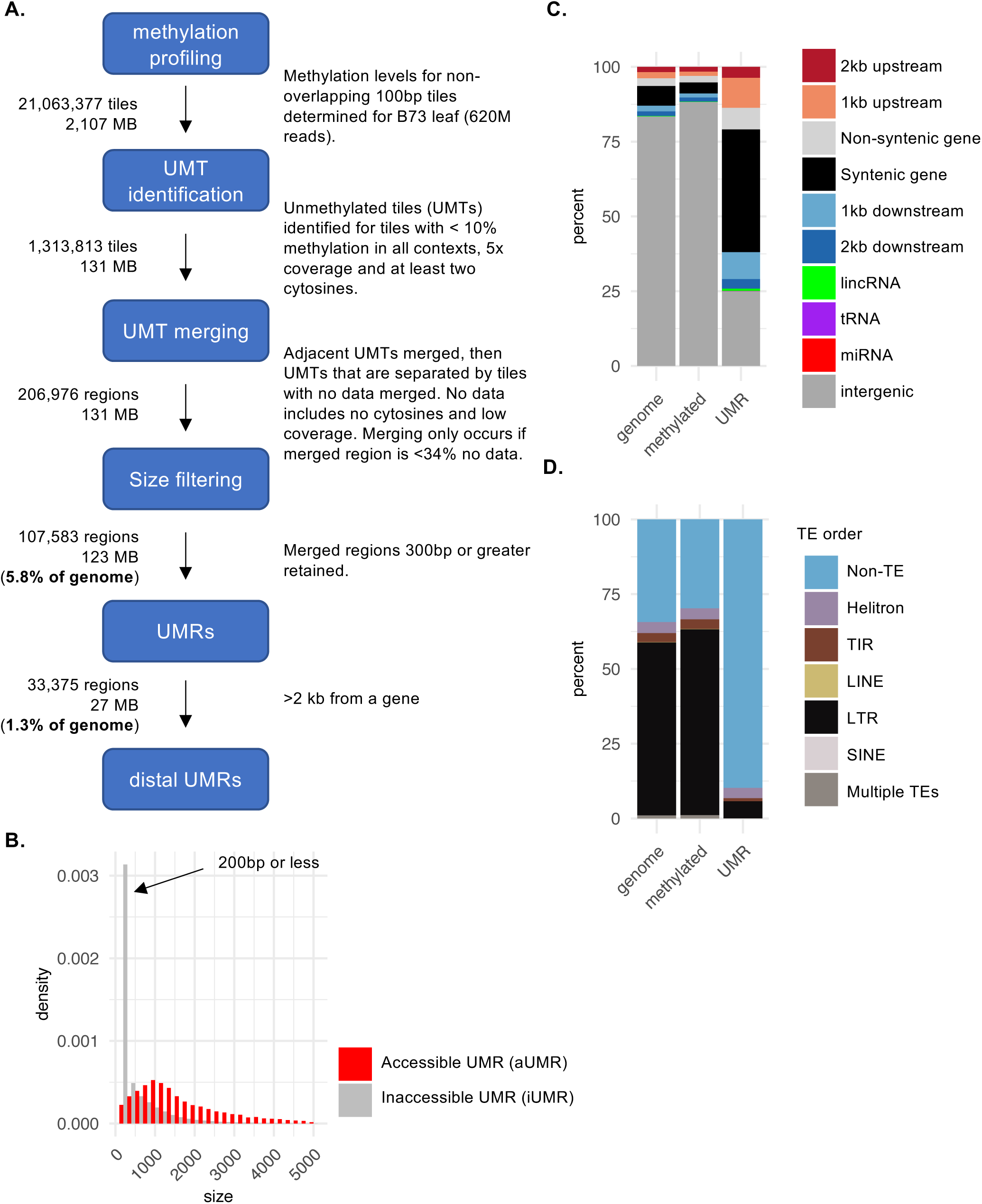
Unmethylated regions in maize. **A**. Pipeline overview of unmethylated region identification in maize leaf. **B**. Size distribution in base pairs of aggregated unmethylated tiles (UMTs) that coincide with accessible chromatin regions (red) compared to aggregated UMTs identified in inaccessible chromatin (grey). The large proportion of inaccessible UMTs are 200 bp or less; therefore these small UMTs are excluded from the final UMR regions. Accessibility determined by overlapping with accessible regions identified in maize leaf in Ricci et al 2019. **C**. Percent of UMTs (100bp tiles) that overlap each genomic feature. Tiles with missing data that are later merged with flanking UMTs are included, all other tiles with missing data or no sites are excluded. **D**. Percent of UMTs (100bp tiles) that overlap transposable elements (TEs). TIR = terminal inverted repeat; LINE = long interspersed nuclear element; LTR = long terminal repeat; SINE = short interspersed nuclear element; 10.7% overlap TEs, including 6.08% LTR, 3.50% Helitron, 1.02% TIR, 0.0388% LINE, 0.0272% SINE and 0.0584 multiple TEs from different orders. Tiles with missing data that are later merged with flanking UMTs are included, while tiles all other tiles with missing data or no sites are excluded.

**Fig S2.**
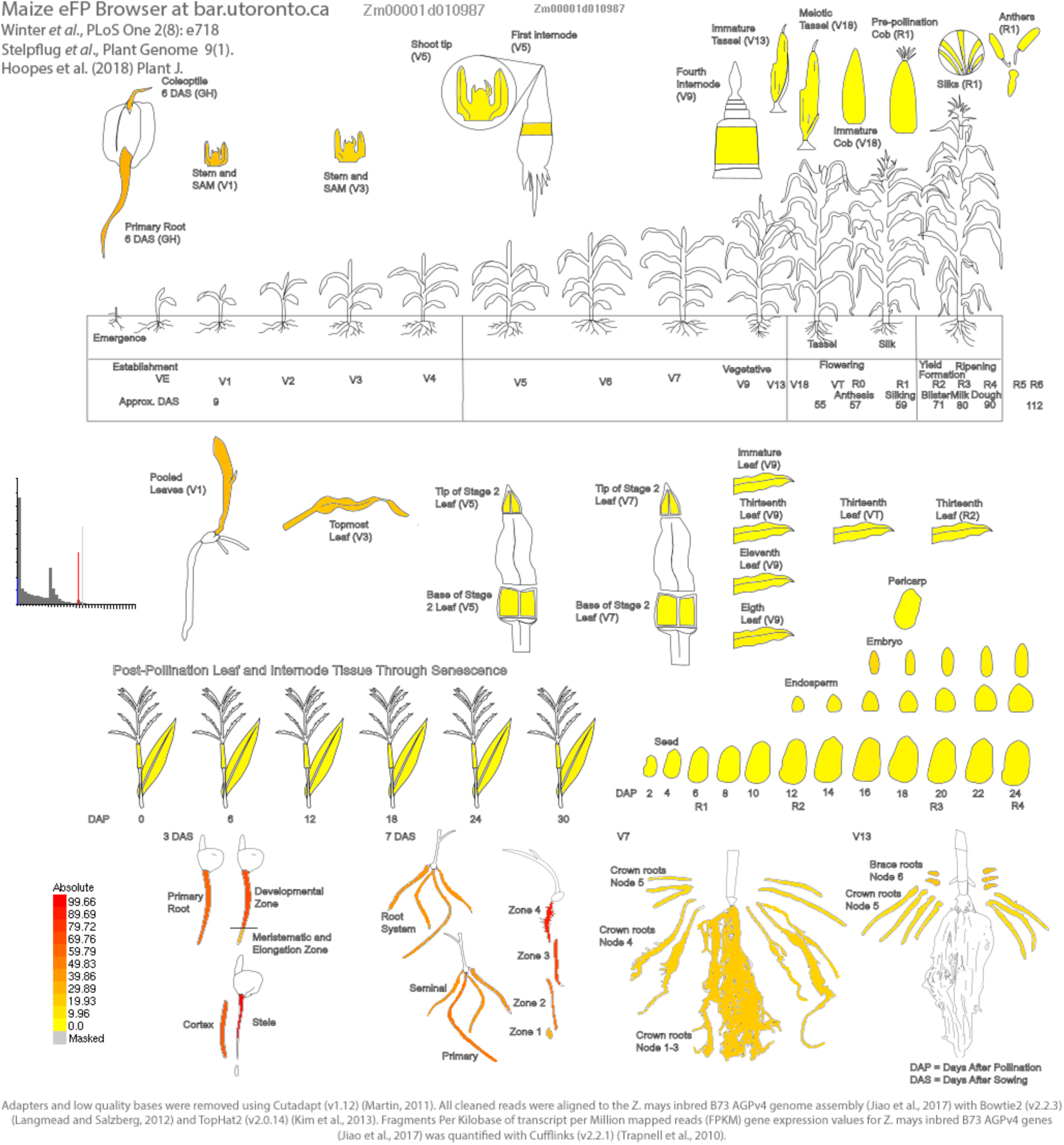
Full expression profile of rap2.7. The eFP browser expression data was downloaded from bar (Winter et al. 2007) hosted on Maize GDB incorporating the maize expression datasets (Stelpflug et al. 2016; Hoopes et al. 2019).

**Fig S3.**
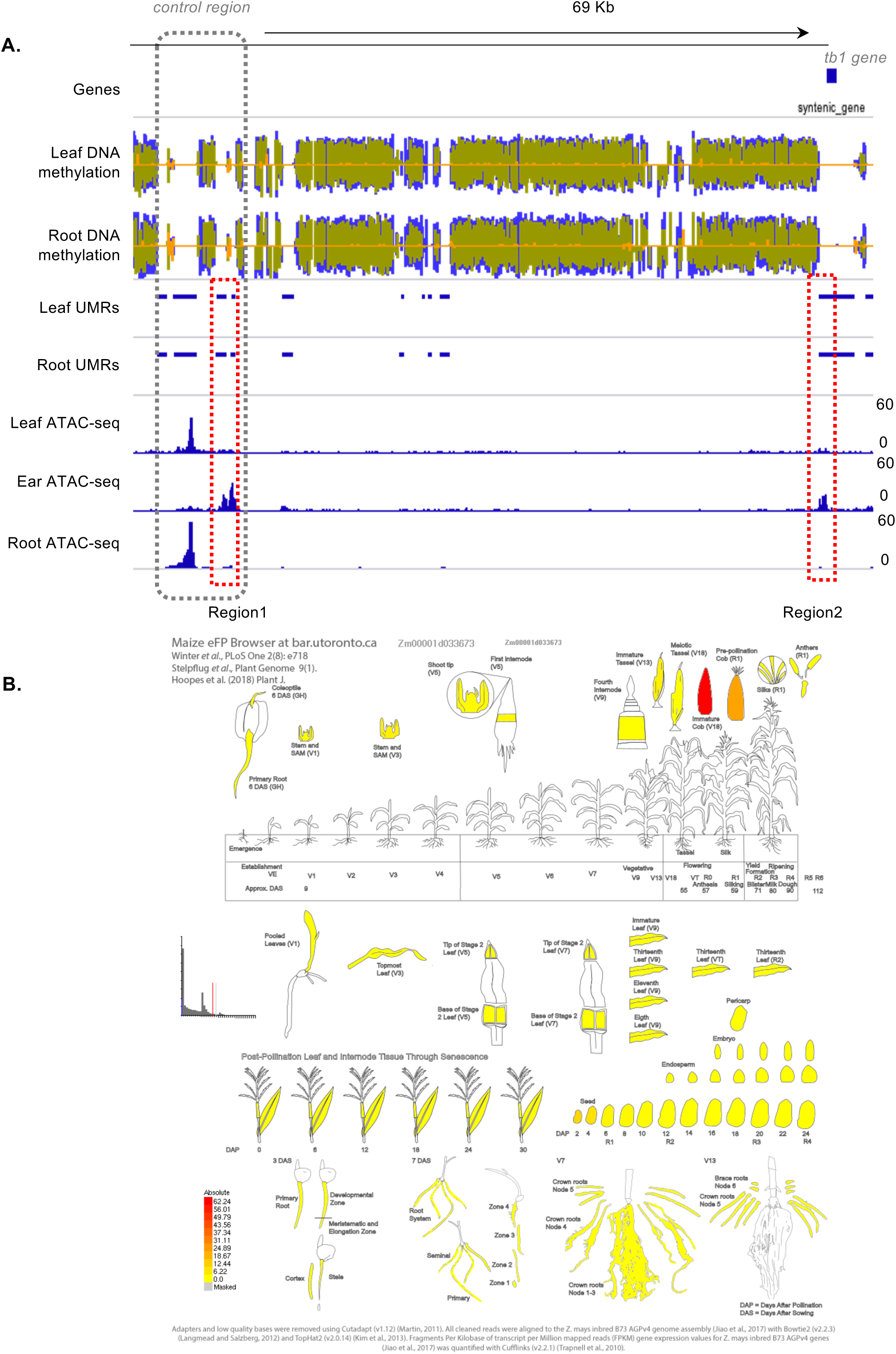
iUMRs that mark tissue-specific accessible regions upstream of the tb1 gene locus. **A**. Example leaf iUMRs that mark regions that become accessible in other tissues. Three **e**xample leaf iUMRs regions in the 70kb genomic region upstream of the tb1 transcription factor. The known cis-regulatory “control” region is marked by the grey dashed box. Region 1 and region 2, marjed by the red dashed boxes, are examples of leaf iUMRs that are inaccessible in leaf but accessible in ear tissue. **B**. The relative gene expression (FPKM) of tb1. The eFP browser expression data was downloaded from bar (Winter et al. 2007) hosted on Maize GDB incorporating the maize expression datasets (Stelpflug et al. 2016; Hoopes et al. 2019).

**Fig S4.**
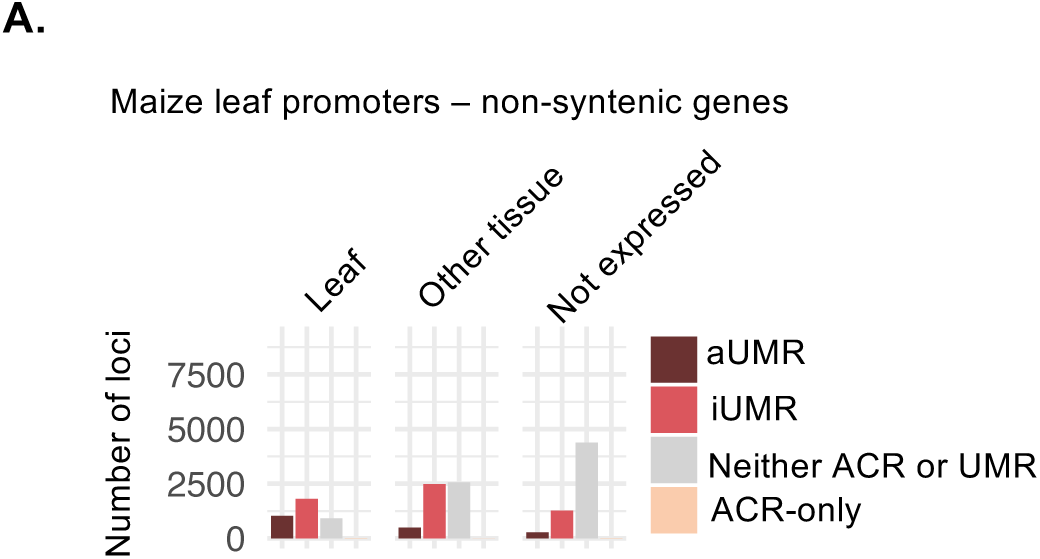
Relationship between expression of a gene and promoter accessibility and methylation for non-syntenic genes. Genes were defined as “leaf” expressed if expression was greater than 1 count per million (CPM) in leaf tissue; “other tissue” expressed if greater than 1 CPM in a tissue other than leaf but less than 1 CPM in leaf; or “not expressed” if expression less than 1 CPM in all tissues.

**Fig S5.**
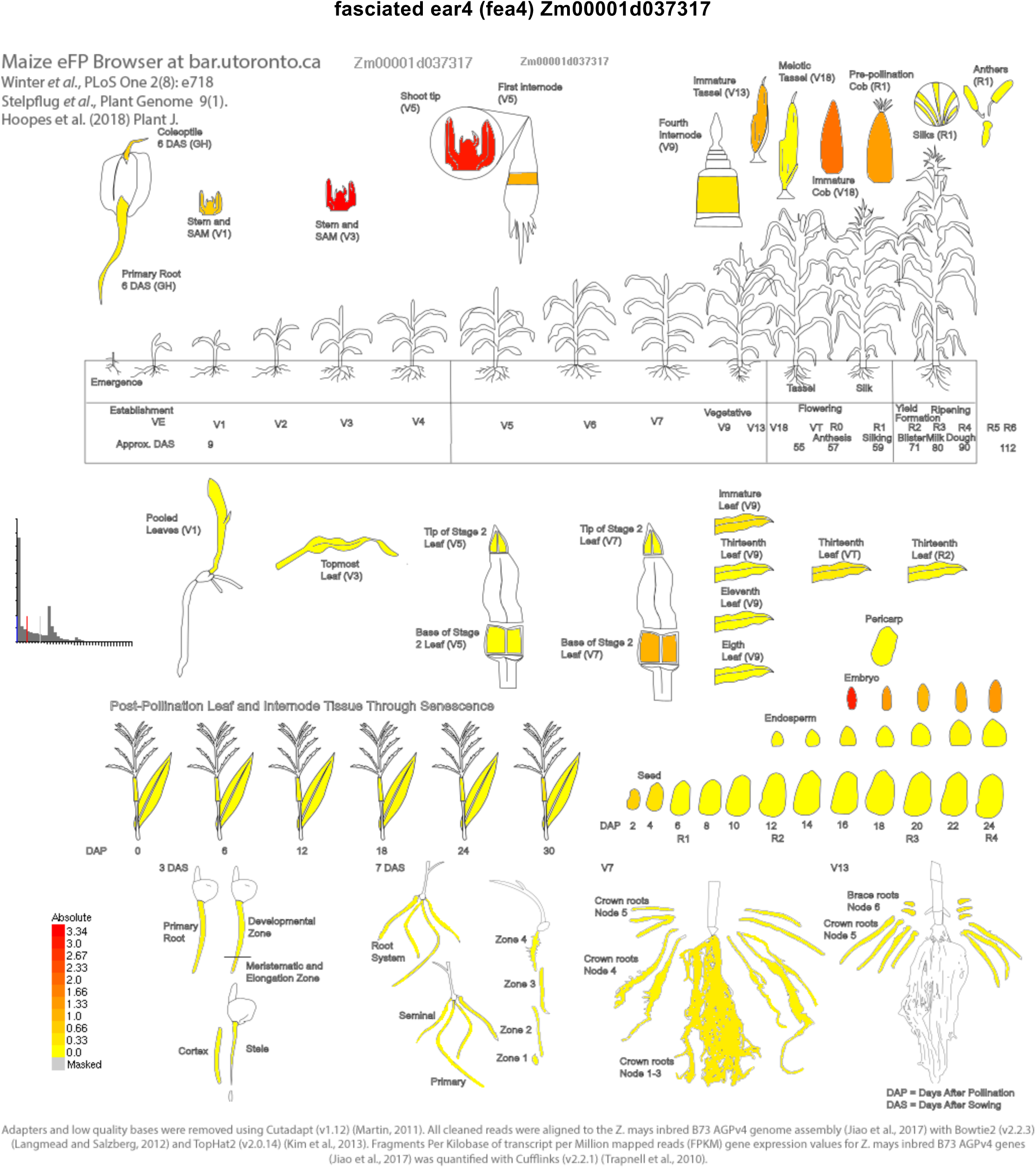
FEA4 expression profile. The eFP browser expression data was downloaded from bar (Winter et al. 2007) hosted on Maize GDB incorporating the maize expression datasets (Stelpflug et al. 2016; Hoopes et al. 2019).

**Fig S6.**
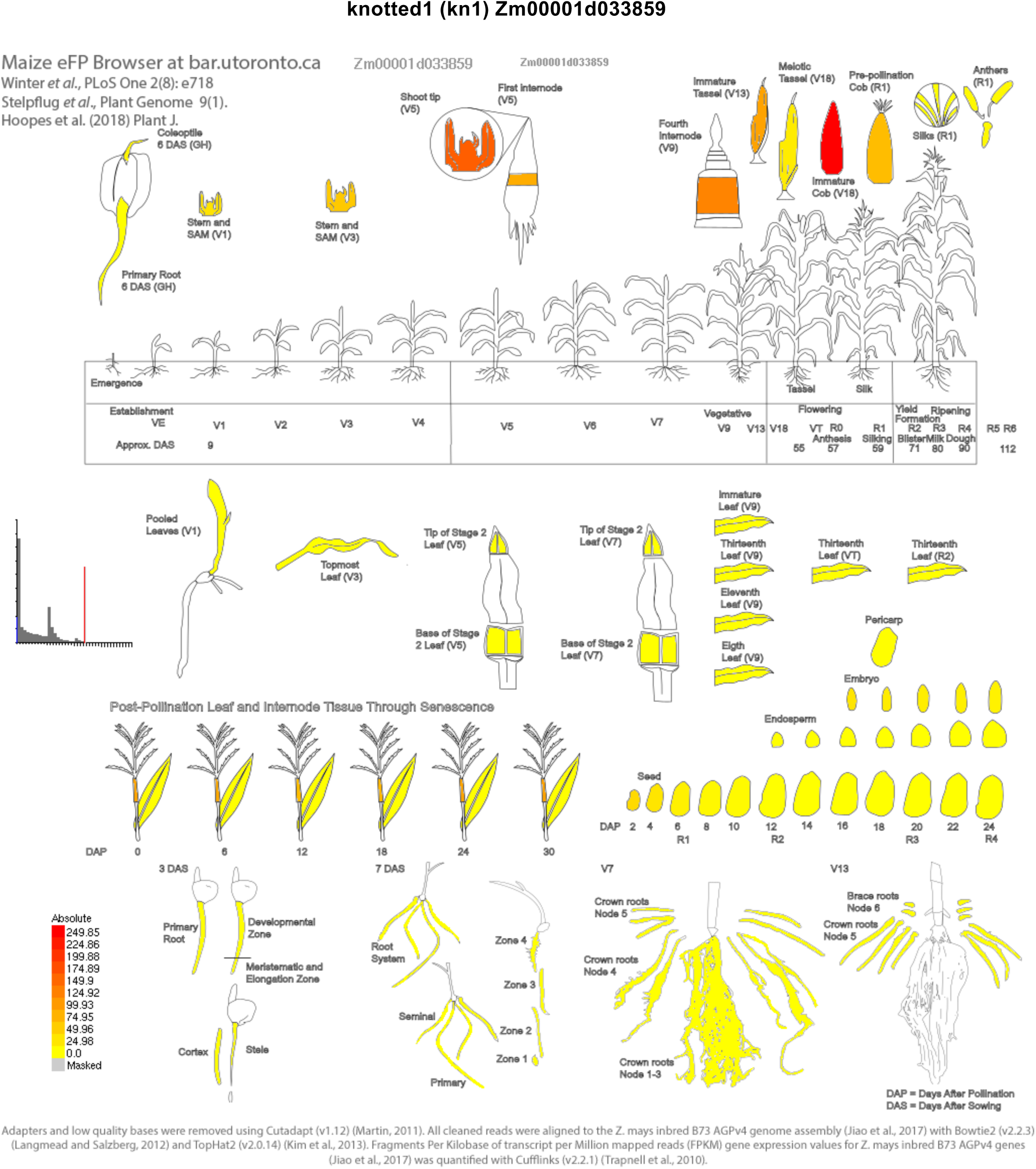
KN1 expression profile. The eFP browser expression data was downloaded from bar (Winter et al. 2007) hosted on Maize GDB incorporating the maize expression datasets (Stelpflug et al. 2016; Hoopes et al. 2019).

**Fig S7.**
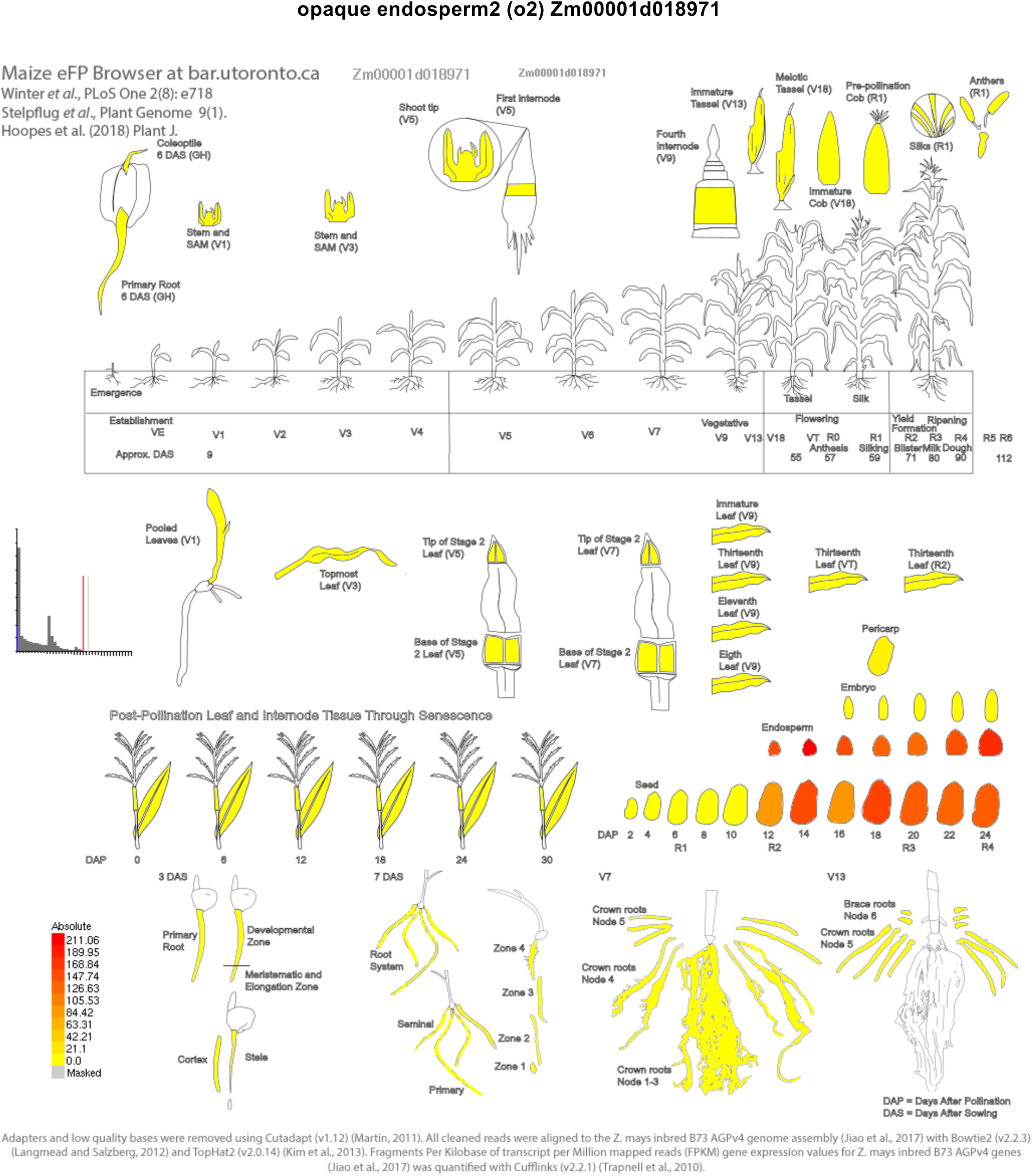
o2 expression profile. The eFP browser expression data was downloaded from bar (Winter et al. 2007) hosted on Maize GDB incorporating the maize expression datasets (Stelpflug et al. 2016; Hoopes et al. 2019).

**Fig S8.**
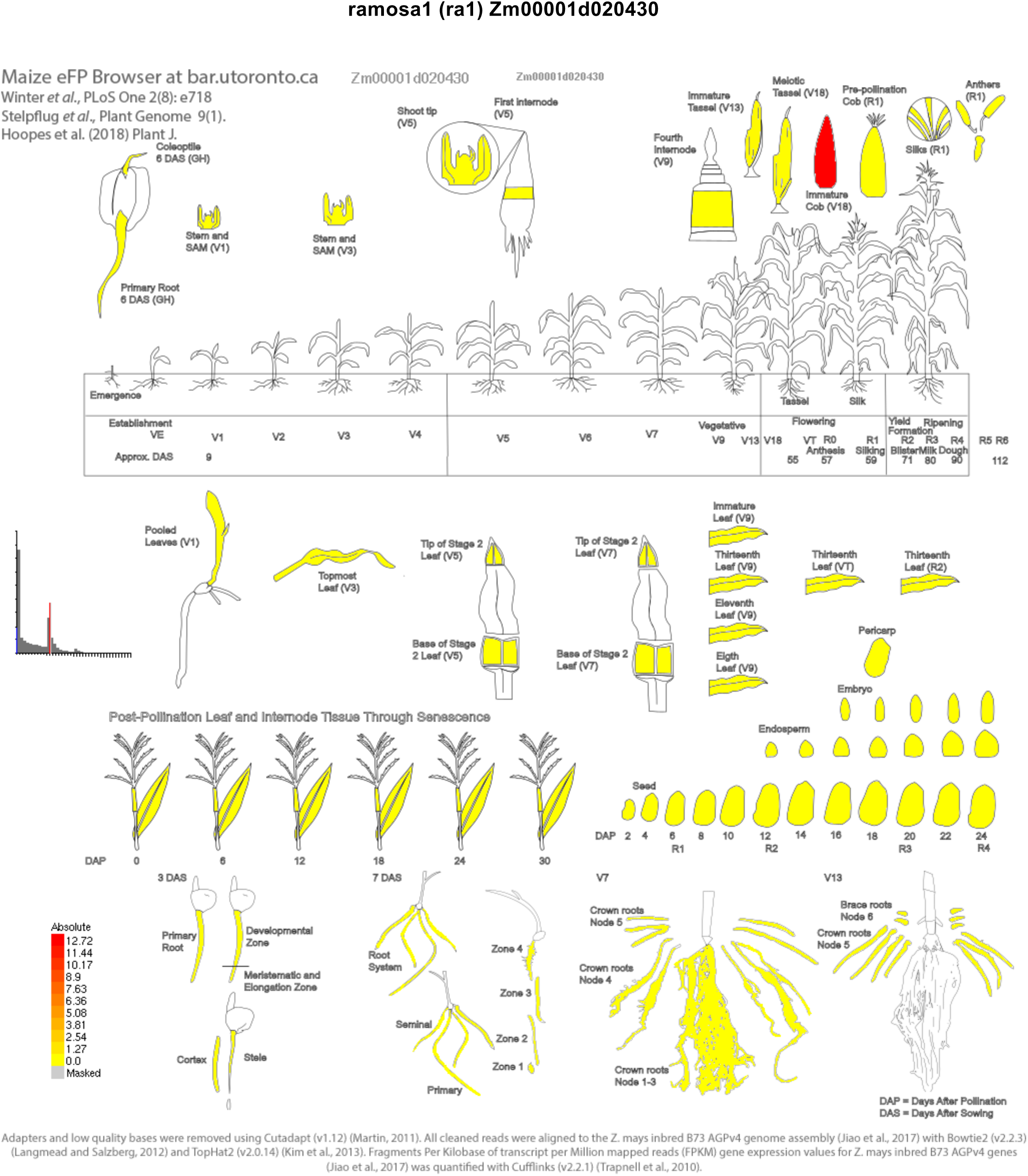
ra1 expression profile. The eFP browser expression data was downloaded from bar (Winter et al. 2007) hosted on Maize GDB incorporating the maize expression datasets (Stelpflug et al. 2016; Hoopes et al. 2019).

**Fig S9.**
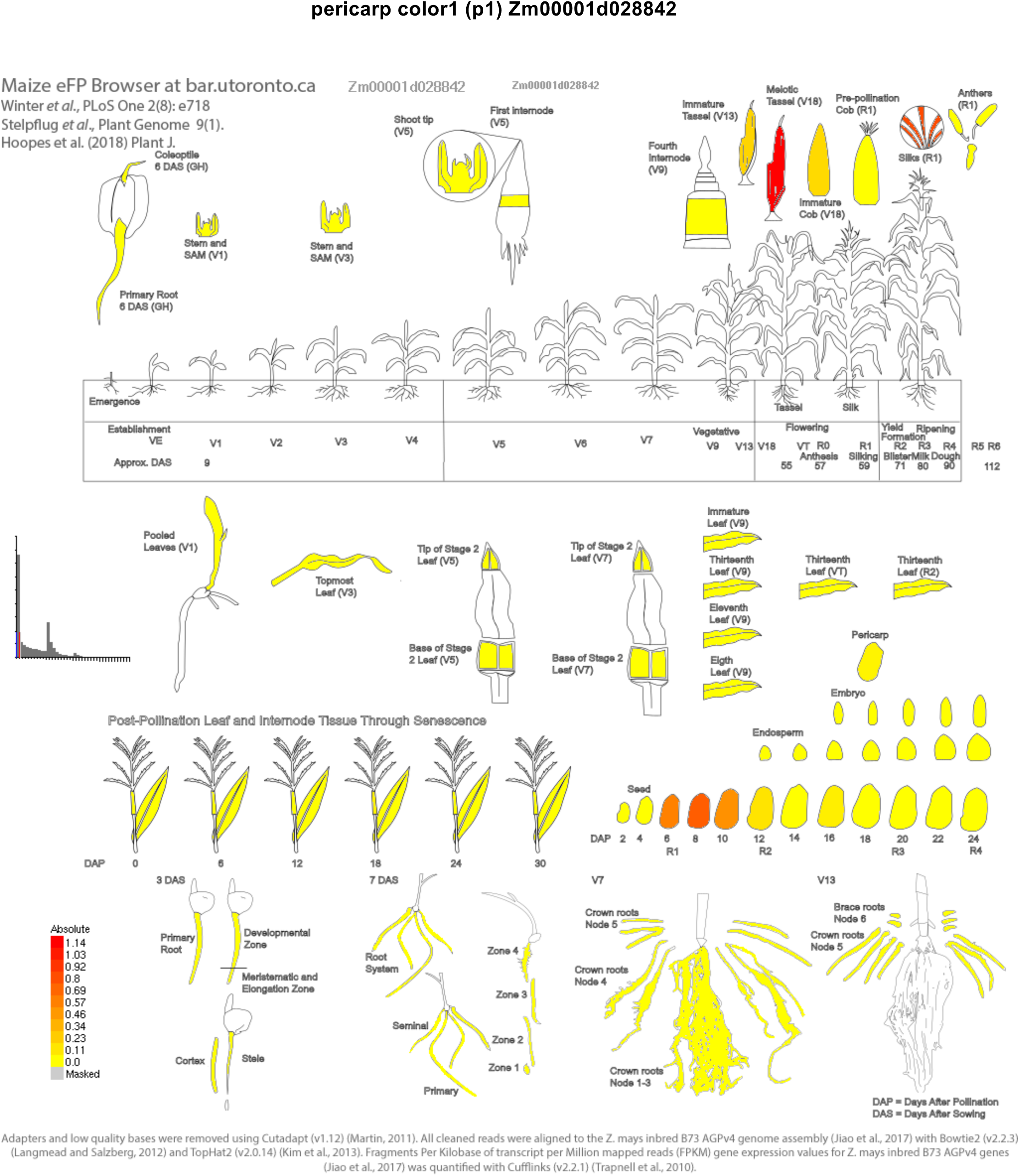
p1 expression profile. The eFP browser expression data was downloaded from bar (Winter et al. 2007) hosted on Maize GDB incorporating the maize expression datasets (Stelpflug et al. 2016; Hoopes et al. 2019).

**Fig S10.**
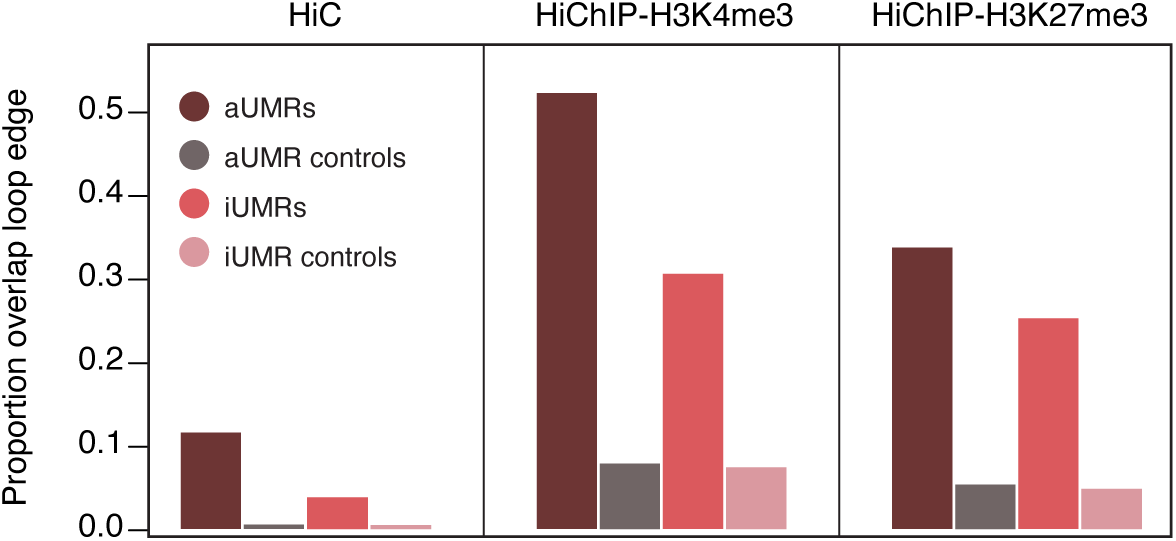
Proportion of aUMRs and iUMRs and associated control regions that overlap at least one loop edge from HiC, H3K4me3-HiChIP and H3K27me3-HiChIP chromatin loops.

